# Genomics of Human Respiratory Syncytial Virus Vaccine Attenuation

**DOI:** 10.1101/862185

**Authors:** Thomas Junier, Laurent Kaiser, Nimisha Chaturvedi, Tina Hartert, Jacques Fellay

## Abstract

The human orthopneumovirus (HRSV) is a major cause of lower respiratory tract infection in children worldwide. Despite decades of efforts, no vaccine is available. In this work, we report mutations that are frequent in vaccine candidates and rare in wild-type genomes, taking into account all the publicly available HRSV sequence data. These mutations are different from the ones already known to attenuate the virus, and thus may contribute to the effort towards producing a live attenuated vaccine against HRSV.

## Introduction

The human orthopneumovirus (known until 2016 as the respiratory syncytial virus, and still referred to as HRSV) is a single-strand, negative-sense RNA virus belonging to family *Pneumoviridae.* It has a 15-kb genome that contains ten genes (Table 1).

**Table 1.** Genes and gene products of the HRSV genome^11^, as well as some of their known functions, by order of position along the (negative-sense) genome. Gene M2 encodes two proteins on different open reading frames on the same mRNA.

| Symbol | Name | Function |
| --- | --- | --- |
| NS1 | Non-structural protein 1 | inhibition of type-I interferon<sup>16</sup> |
| NS2 | Non-structural protein 1B | inhibition of RIG-I<sup>17</sup> |
| N | Nucleoprotein | encapsidation<sup>18</sup> |
| P | Phosphoprotein | stabilization of the polymerase<sup>19</sup> |
| M | Matrix protein | virion assembly<sup>20</sup> |
| SH | Small hydrophobic protein | ion transport<sup>21</sup> |
| G | Major surface glycoprotein G | attachment to host cell<sup>22</sup> |
| F | Fusion glycoprotein F0 | fusion of viral and cell membranes<sup>23</sup> |
| M2 | Matrix M2 | transcription<sup>24</sup> |
| L | RNA-directed RNA polymerase L | replication, transcription<sup>25</sup> |

HRSV infects the respiratory tract, and causes a variety of symptoms ranging from mild to fatal. It has very high lifetime prevalence: almost everyone is infected for the first time by age two. It is a major cause of acute lower respiratory tract infection in children worldwide, and caused an estimated 50,000 to 75,000 in-hospital deaths in children under five years old around the world in 2015, the vast majority in developing countries^1, 2^. HRSV thus ranks among the leading causes of death in children^3^. In the USA, HRSV causes 100-500 yearly deaths in children under five years old (as of 2020), while the number of deaths among adults is around 14,000^4^. With an estimated 19.6 million children under 5 and 254 million adults (over 18) in the USA^5^, the mortality rates are around 25 per million and 55 per million, respectively. Among adults, the elderly are most at risk, both in terms of incidence and severity.^6^.

HRSV is divided into two subtypes, A and B. Subtype A appears to be more frequent in infected patients.^7^ Re-infections can occur at any age, but symptoms are most severe in children under one year old, the elderly, and immunocompromised patients.^8^ While prophylaxis (a monoclonal antibody marketed as Palivizumab®) and treatment (the guanosine analog Ribavirin®) against HRSV exist, no vaccine is currently available. Palivizumab and Ribavirin are expensive (the latter is also toxic^9^) and only recommended for high-risk patients^10^, which makes the development of a safe and effective vaccine a high priority.

In the 1960s, an early attempt at a HRSV vaccine using formalin-inactivated virus ended in failure: the vaccine did not protect against infection and in some cases actually caused enhanced disease (including fatalities).^11^ Inactivated vaccines against HRSV are no longer pursued since this setback. Modern approaches include particulate or subunit vaccines, gene-based vaccines, and live attenuated vaccines (LAVs).

Live attenuated vaccines are pathogens that are able to replicate in the patient, but are less virulent (“attenuated”) than the wild type. LAVs have been developed against measles, yellow fever, smallpox and polio, among others; they were instrumental in the eradication or near-eradication of the last two.

One method for achieving attenuation involves several generations of culture in an environment different from the original host cell (for example, a different kind of cell or a lower temperature (“cold passage”)). As the virus adapts to its new environment, it may become less adapted to the original one - thus losing virulence - while still being able to elicit a strong immune response from the host. Another approach involves chemical mutagenesis followed by selection for temperature-sensitive mutants.

LAVs offer some advantages, including the fact that they do not cause enhanced disease, that they stimulate immunity both systemically and in the respiratory tract, that they can be delivered intranasally, and that they replicate in young infants even in the presence of maternal HRSV antibody.^12, 13^ They also do not require adjuvants^14^. LAVs are not without drawbacks, however: their use might be limited to children under two years^13^, and they may pose some risks, especially in immunocompromised patients and children^15^,

Understanding attenuating mutations is useful for at least the following reasons: first, attenuation achieved by a smaller number of mutations has a higher risk of reversion to virulence than attenuation achieved by a larger number; second, a knowledge of the phenotypic effect of mutations enables a reverse genetics approach to LAV production, in which desired combinations of attenuating mutations are deliberately introduced into the viral genome; finally, the effect of mutations is not always additive: some mutations may in fact even cancel one another - for example, one mutation in the L gene is known to compensate another mutation in the same gene^12^ - hence the importance of studying the effects of combinations of mutations. The hope is, in the end, that a better understanding of HRSV genomics may help overcome the challenge posed by producing a safe yet suffciently immunogenic vaccine^13^.

Attenuating mutations have been identified by Karron *et al.* through whole-genome sequencing of a cold-passaged strain as well as six temperature-sensitive derivatives of that strain.^12^. In this work, we followed a comparable approach, but using all publicly available HRSV sequence data: the public sequence databases contain thousands of HRSV sequences, both wild-type and attenuated, which we aligned together then scanned for positions in which the two groups have different allele frequencies.

## Results

### Breakdown of Wild-type Sequences by Subtype and Gene

Out of the 28,248 GenBank entries originally downloaded, 11,600 were rejected due to being of unknown or explicitly non-HRSV origin, leaving 16,648. Among these, inference of category (vaccine vs. wild-type) recognized 575 vaccine and 16,073 wild-type entries. Filtering the latter by quality, followed by inference of subtype and gene and finally by extraction of coding sequences (CDSs) yielded 10,975 wild-type sequences, of which a breakdown is shown in table 2, column 3.

**Table 2.**
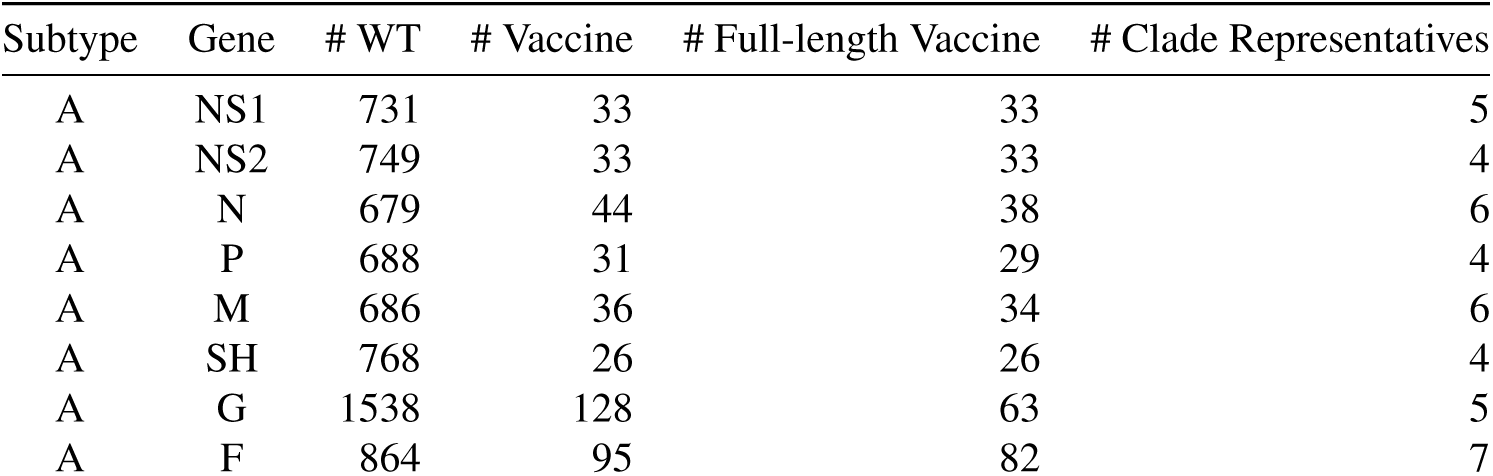
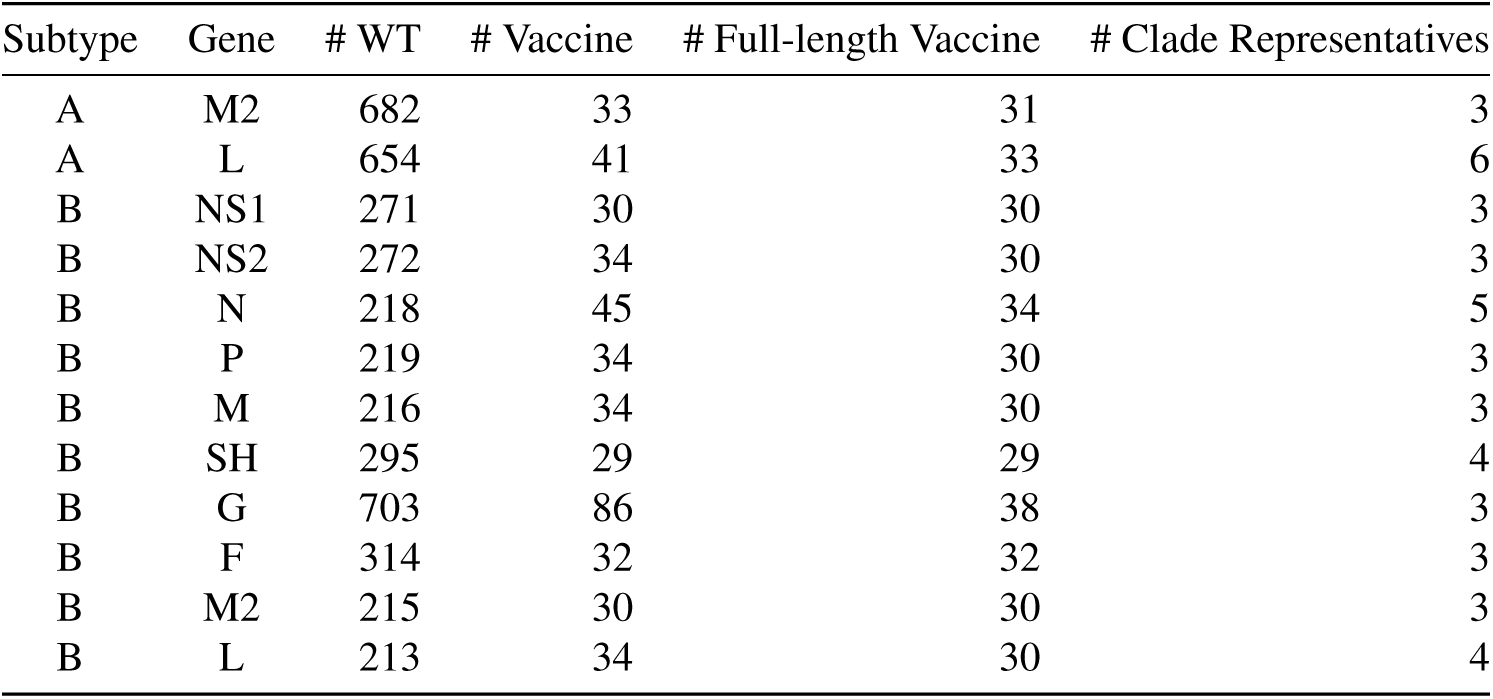
Breakdown of wild-type and vaccine sequences by HRSV subtype and gene.

| Subtype | Gene | # WT | # Vaccine | # Full-length Vaccine | # Clade Representatives |
| --- | --- | --- | --- | --- | --- |
| A | NS1 | 731 | 33 | 33 | 5 |
| A | NS2 | 749 | 33 | 33 | 4 |
| A | N | 679 | 44 | 38 | 6 |
| A | P | 688 | 31 | 29 | 4 |
| A | M | 686 | 36 | 34 | 6 |
| A | SH | 768 | 26 | 26 | 4 |
| A | G | 1538 | 128 | 63 | 5 |
| A | F | 864 | 95 | 82 | 7 |

| Subtype | Gene | # WT | # Vaccine | # Full-length Vaccine | # Clade Representatives |
| --- | --- | --- | --- | --- | --- |
| A | M2 | 682 | 33 | 31 | 3 |
| A | L | 654 | 41 | 33 | 6 |
| B | NS1 | 271 | 30 | 30 | 3 |
| B | NS2 | 272 | 34 | 30 | 3 |
| B | N | 218 | 45 | 34 | 5 |
| B | P | 219 | 34 | 30 | 3 |
| B | M | 216 | 34 | 30 | 3 |
| B | SH | 295 | 29 | 29 | 4 |
| B | G | 703 | 86 | 38 | 3 |
| B | F | 314 | 32 | 32 | 3 |
| B | M2 | 215 | 30 | 30 | 3 |
| B | L | 213 | 34 | 30 | 4 |

### Classification of Vaccine Sequences by Subtype and Gene

Column 4 of table 2 shows a breakdown of vaccine sequences (or sub-sequences, in the case of sequences matching more than one gene) by subtype and gene. Column 5 shows the number of sequences for each (subtype,gene) combination after discarding sequences that were < 90% full length, as was done for wild-type sequences.

### Gene Phylogenies

Eevery gene except F shows a clear phylogenetic separation of the two subtypes (Figure 1, see supplementary data for all trees in Newick format and as graphical representations). It also shows that vaccines of either subtype do not form a single clade, but are instead spread out over the whole respective A or B subtree. Finally, each tree shows vaccine sequences arranged in a distinctive, stair-like pattern consisting of wholly unbalanced nodes, that is, nodes of which one child is a leaf and all other descendants descend from the other child, which is itself wholly unbalanced (Figure 2).

**Figure 1.**
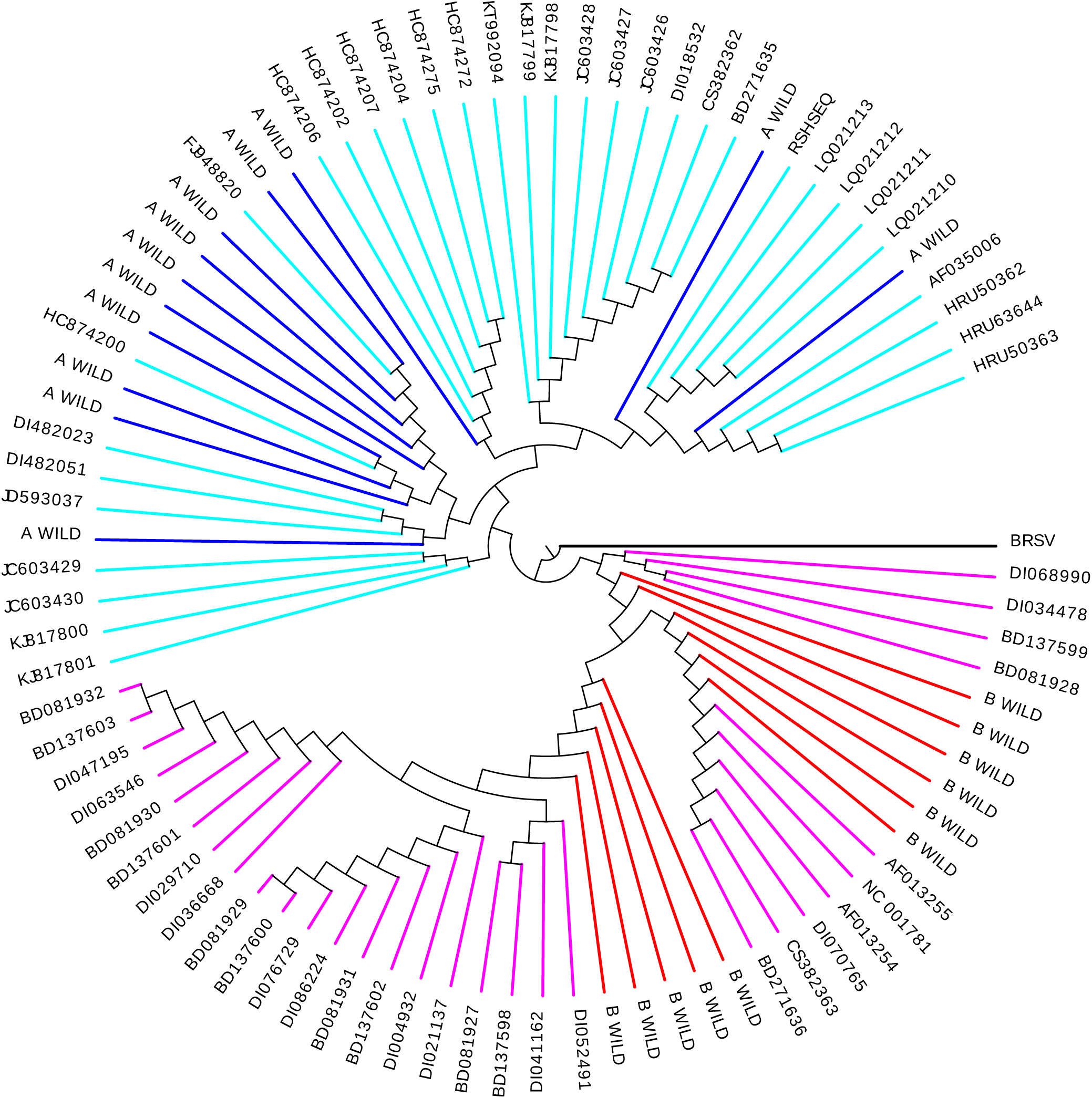
The phylogeny of gene L. Subtype A is in blue (wild-type) and cyan (vaccines); subtype B in red (wild-type) and magenta (vaccines). The tree was re-rooted on the bovine HRSV ortholog, the purely vaccine clades condensed into single leaves, and the resulting tree rendered as SVG using the Newick Utilities.^26^

**Figure 2.**
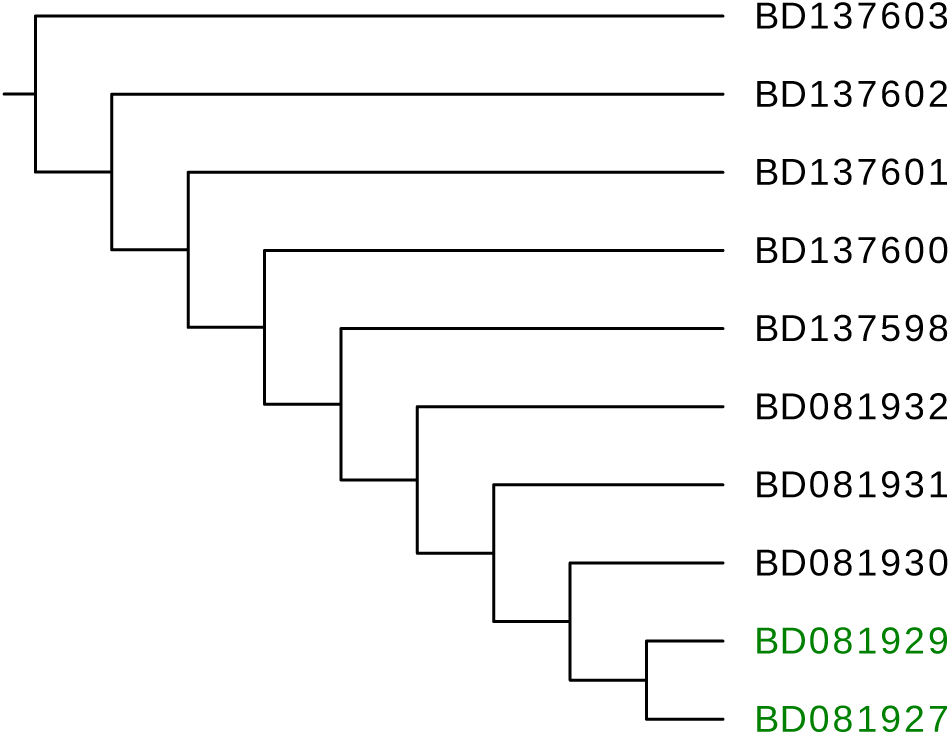
An extremely unbalanced clade. This is a subtree of the F gene tree. From such a clade we only retain one (at random) of the “end-of-chain” sequences, marked in green. These are likely to contain at least as many mutations as any other sequence in the clade.

### Clade Representatives

The selection of one vaccine per unbalanced clade further reduced the number of vaccine sequences (Table 2, column 6). Pooled with the corresponding wild-type sequences, they yielded twenty sets of sequences, one each per (gene, subtype) pair, consisting only of full-length or nearly full-length sequences, and with a minimal number of close relatives among the vaccines.

### Vaccine-linked Mutations

Table 3 shows all vaccine-linked, non-silent mutations for which the Fisher exact test’s *p*-value is not greater than the significance threshold *α* = 0.01, after applying the Bonferroni correction for 6573 simultaneous comparisons (one per non-conserved position in the genome) - this results in a corrected *α* = 1.52 × 10^−6^. No position matching these criteria was found in subtype B. Column 12 of the table shows the effect size as the odds ratio of a vaccine vs. a wild-type sequence having the minor allele at a given position, flanked by the lower bound of the 95% confidence interval around the ratio (column 12) and the upper bound (column 13). None of the intervals includes the value 1, indeed the minimal lower bound is 12.

**Table 3.**
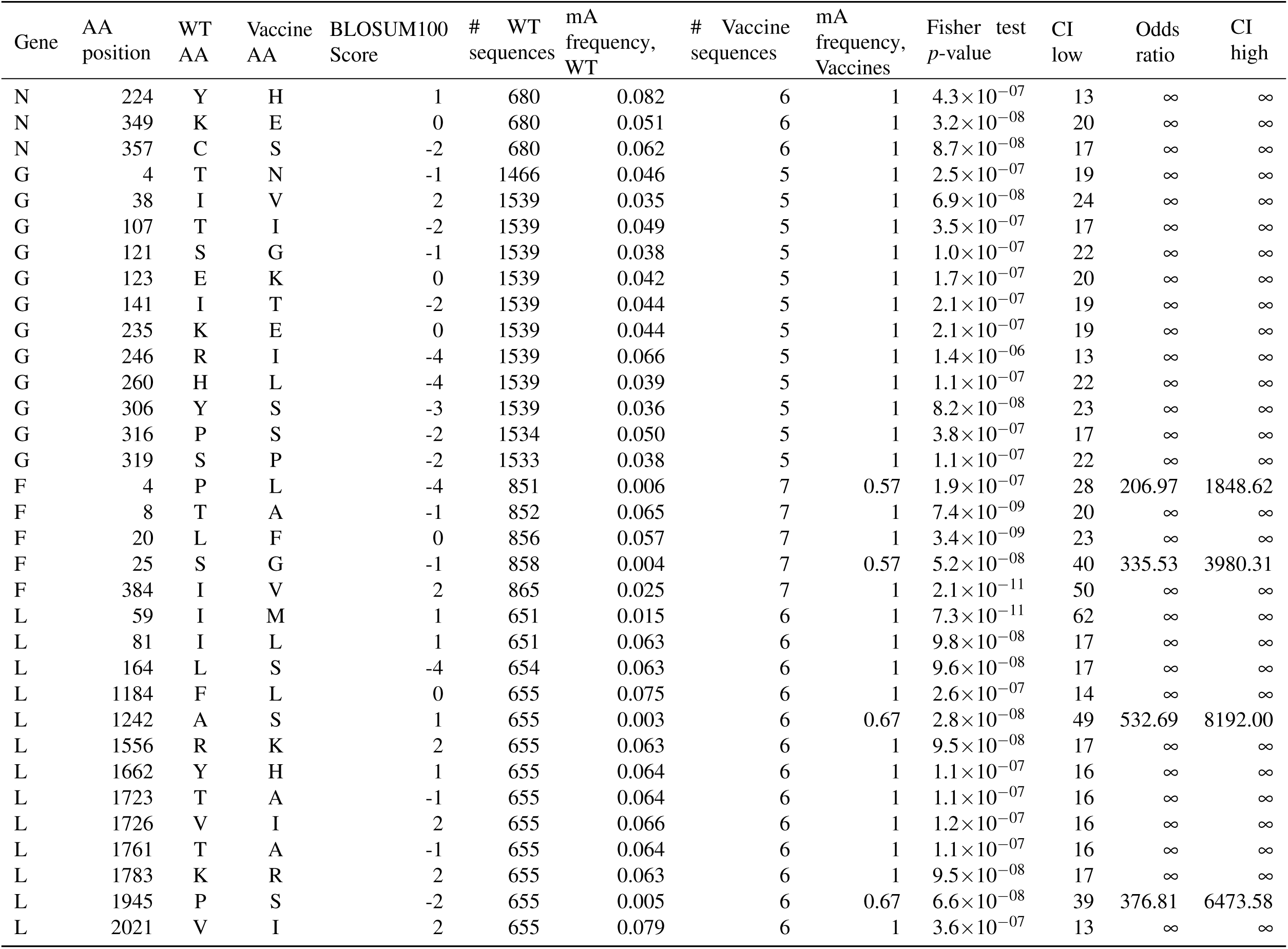
Vaccine-linked, non-silent mutations. The amino acid (AA) position is with reference to `GU591769`; the wild-type (WT) and vaccine amino acids columns list the most frequent amino acid in wild-type and vaccine sequences, respectively. Fisher’s test compares minor allele (mA) frequencies for vaccines and WT, given the counts of sequences in each. Only positions with p-value lower than *α* = 0.01 corrected for multiple comparisons are shown. No such positions were found in subtype B. The effect size is given as the odds ratio between mA frequencies in vaccines versus WT. CI: 95% confidence interval for the odds ratio.

| Gene | AA position | WT AA | Vaccine AA | BLOSUM100 Score | # WT sequences | mA frequency, WT | # Vaccine sequences | mA frequency, Vaccines | Fisher test $p$-value | CI low | Odds ratio | CI high |
| --- | --- | --- | --- | --- | --- | --- | --- | --- | --- | --- | --- | --- |
| N | 224 | Y | H | 1 | 680 | 0.082 | 6 | 1 | $4.3 \times 10^{-07}$ | 13 | ∞ | ∞ |
| N | 349 | K | E | 0 | 680 | 0.051 | 6 | 1 | $3.2 \times 10^{-08}$ | 20 | ∞ | ∞ |
| N | 357 | C | S | -2 | 680 | 0.062 | 6 | 1 | $8.7 \times 10^{-08}$ | 17 | ∞ | ∞ |
| G | 4 | T | N | -1 | 1466 | 0.046 | 5 | 1 | $2.5 \times 10^{-07}$ | 19 | ∞ | ∞ |
| G | 38 | I | V | 2 | 1539 | 0.035 | 5 | 1 | $6.9 \times 10^{-08}$ | 24 | ∞ | ∞ |
| G | 107 | T | I | -2 | 1539 | 0.049 | 5 | 1 | $3.5 \times 10^{-07}$ | 17 | ∞ | ∞ |
| G | 121 | S | G | -1 | 1539 | 0.038 | 5 | 1 | $1.0 \times 10^{-07}$ | 22 | ∞ | ∞ |
| G | 123 | E | K | 0 | 1539 | 0.042 | 5 | 1 | $1.7 \times 10^{-07}$ | 20 | ∞ | ∞ |
| G | 141 | I | T | -2 | 1539 | 0.044 | 5 | 1 | $2.1 \times 10^{-07}$ | 19 | ∞ | ∞ |
| G | 235 | K | E | 0 | 1539 | 0.044 | 5 | 1 | $2.1 \times 10^{-07}$ | 19 | ∞ | ∞ |
| G | 246 | R | I | -4 | 1539 | 0.066 | 5 | 1 | $1.4 \times 10^{-06}$ | 13 | ∞ | ∞ |
| G | 260 | H | L | -4 | 1539 | 0.039 | 5 | 1 | $1.1 \times 10^{-07}$ | 22 | ∞ | ∞ |
| G | 306 | Y | S | -3 | 1539 | 0.036 | 5 | 1 | $8.2 \times 10^{-08}$ | 23 | ∞ | ∞ |
| G | 316 | P | S | -2 | 1534 | 0.050 | 5 | 1 | $3.8 \times 10^{-07}$ | 17 | ∞ | ∞ |
| G | 319 | S | P | -2 | 1533 | 0.038 | 5 | 1 | $1.1 \times 10^{-07}$ | 22 | ∞ | ∞ |
| F | 4 | P | L | -4 | 851 | 0.006 | 7 | 0.57 | $1.9 \times 10^{-07}$ | 28 | 206.97 | 1848.62 |
| F | 8 | T | A | -1 | 852 | 0.065 | 7 | 1 | $7.4 \times 10^{-09}$ | 20 | ∞ | ∞ |
| F | 20 | L | F | 0 | 856 | 0.057 | 7 | 1 | $3.4 \times 10^{-09}$ | 23 | ∞ | ∞ |
| F | 25 | S | G | -1 | 858 | 0.004 | 7 | 0.57 | $5.2 \times 10^{-08}$ | 40 | 335.53 | 3980.31 |
| F | 384 | I | V | 2 | 865 | 0.025 | 7 | 1 | $2.1 \times 10^{-11}$ | 50 | ∞ | ∞ |
| L | 59 | I | M | 1 | 651 | 0.015 | 6 | 1 | $7.3 \times 10^{-11}$ | 62 | ∞ | ∞ |
| L | 81 | I | L | 1 | 651 | 0.063 | 6 | 1 | $9.8 \times 10^{-08}$ | 17 | ∞ | ∞ |
| L | 164 | L | S | -4 | 654 | 0.063 | 6 | 1 | $9.6 \times 10^{-08}$ | 17 | ∞ | ∞ |
| L | 1184 | F | L | 0 | 655 | 0.075 | 6 | 1 | $2.6 \times 10^{-07}$ | 14 | ∞ | ∞ |
| L | 1242 | A | S | 1 | 655 | 0.003 | 6 | 0.67 | $2.8 \times 10^{-08}$ | 49 | 532.69 | 8192.00 |
| L | 1556 | R | K | 2 | 655 | 0.063 | 6 | 1 | $9.5 \times 10^{-08}$ | 17 | ∞ | ∞ |
| L | 1662 | Y | H | 1 | 655 | 0.064 | 6 | 1 | $1.1 \times 10^{-07}$ | 16 | ∞ | ∞ |
| L | 1723 | T | A | -1 | 655 | 0.064 | 6 | 1 | $1.1 \times 10^{-07}$ | 16 | ∞ | ∞ |
| L | 1726 | V | I | 2 | 655 | 0.066 | 6 | 1 | $1.2 \times 10^{-07}$ | 16 | ∞ | ∞ |
| L | 1761 | T | A | -1 | 655 | 0.064 | 6 | 1 | $1.1 \times 10^{-07}$ | 16 | ∞ | ∞ |
| L | 1783 | K | R | 2 | 655 | 0.063 | 6 | 1 | $9.5 \times 10^{-08}$ | 17 | ∞ | ∞ |
| L | 1945 | P | S | -2 | 655 | 0.005 | 6 | 0.67 | $6.6 \times 10^{-08}$ | 39 | 376.81 | 6473.58 |
| L | 2021 | V | I | 2 | 655 | 0.079 | 6 | 1 | $3.6 \times 10^{-07}$ | 13 | ∞ | ∞ |

### Number of mutations in vaccine versus wild-type sequences

Vaccine sequences were heavily mutated at the vaccine-linked positions (Table 4). Indeed, in the attachment glycoprotein and nucleoprotein, *every* vaccine sequence differs from the wild type at *all* such positions, and even in the fusion glycoprotein and polymerase the average number of mutated positions per sequence (4.14 and 12.33) are close to the maxima (5 and 13), respectively.

**Table 4.** Number of mutations in vaccine vs. wild-type sequences (subtype A).

| Gene | # mutated positions | # WT sequences | # WT mutations | avg # mutations / WT sequence | # vaccine sequences | # vaccine mutations | avg # mutations / vaccine sequences | avg mutation ratio |
| --- | --- | --- | --- | --- | --- | --- | --- | --- |
| N | 3 | 679 | 133 | 0.20 | 6 | 18 | 3.00 | 15.32 |
| G | 11 | 1538 | 1488 | 0.97 | 5 | 55 | 11.00 | 11.37 |
| F | 5 | 864 | 188 | 0.22 | 7 | 29 | 4.14 | 19.04 |
| L | 13 | 654 | 753 | 1.15 | 6 | 74 | 12.33 | 10.71 |

## Discussion

### Power Considerations

The fact that very few vaccine sequences are retained raises the question of the power of our statistical procedure. However, estimates (using R’s `power.fisher.test` function, from package `statmod`) show that even with fewer than ten vaccines a power higher than 0.8 can be achieved if the allele frequencies are sufficiently different. For example, consider a position in which only 5 out of 851 wild-type sequences have the minor allele while 4 out of 7 vaccine sequences have it (as is the case of gene F for subtype A at nucleotide position 11), corresponding to relative frequencies of 0.006 and 0.57, respectively: power estimates (with *α =* 0.01 and 1,000 simulations) consistently exceed 0.95.

### Selection Pressure

Any position under positive selection pressure will exhibit a change in allele frequency, hence it is likely to belong to the set of positions presented in table 3. The converse is generally not true, however, therefore it cannot be inferred from the presence of a position in table 3 that it must be under selection pressure.

### Non-attenuated Vaccines

Vaccine sequences not explicitly annotated as “attenuated” may result from a process other than attenuation and thus not harbour attenuation-linked mutations. This will result in an underestimation of the frequency of these mutations among attenuated sequences, resulting in a higher false negative rate.

### Independence of Mutation events

The phylogenetic trees constructed for each gene of both HRSV subtypes clearly show extremely unbalanced nodes, that is, nodes of which one child is a single leaf, and the other child has several descendants. That other child node is frequently itself extremely unbalanced, resulting in strongly or even wholly unbalanced clades (Figure 2).

Such clade topologies are expected to arise when a sample is taken from a culture to start a new culture, and the process is iteratively repeated: all cultures but the first are more closely related to each other than any of them is to the first culture, and the same situation obtains recursively *within* the last-but-one cultures. The result is an extremely unbalanced phylogeny.

Any early mutation will therefore (barring reversion) be inherited by most of the members of the clade, and be found at a high frequency. However, this high frequency is not indicative of a high number of independent mutation events, but rather, of inheritance. It is therefore wrong to interpret this high frequency as a sign of positive selection. In the context of this work, the mutation would be wrongly thought of as a result of the attenuation process.

Our approach to this problem, namely the selection of a single sequence per fully unbalanced vaccine clade, makes it less likely that any similarities are due to shared ancestry. Furthermore, the fact that we select among the sequences with the maximal number of ancestors in the clade should maximize the number of mutations that we are able to detect.

### Effect on Phenotype

Most mutations retained by our pipeline are silent. Among the non-silent ones, the most likely to affect the phenotype are arguably those which are most rarely encountered between homologous sequences, as measured for example by the BLOSUM^27^ amino acid substitution scores.

Another approach to estimating the effect of a mutation is to consider any known function of the affected protein or regions therein. Table 5 shows annotations pertaining to regions harbouring mutations, according to UniProtKB^28^. Mutations at these sites can thus at least conceivably impair the corresponding stages of the viral cycle.

**Table 5.**
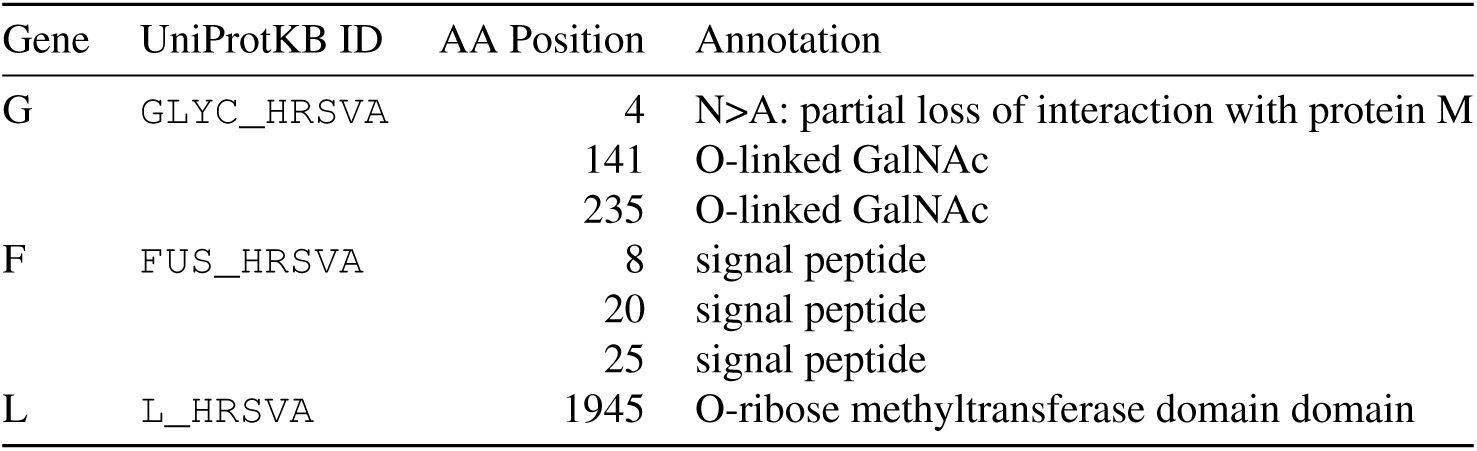
Annotations in UniProtKB sequences for regions/positions corresponding to vaccine mutations.

| Gene | UniProtKB ID | AA Position | Annotation |
| --- | --- | --- | --- |
| G | GLYC_HRSVA | 4 | N>A: partial loss of interaction with protein M |
|  |  | 141 | O-linked GalNAc |
|  |  | 235 | O-linked GalNAc |
| F | FUS_HRSVA | 8 | signal peptide |
|  |  | 20 | signal peptide |
|  |  | 25 | signal peptide |
| L | L_HRSVA | 1945 | O-ribose methyltransferase domain domain |

### Known Attenuation Variants

UniProtKB entry `FUS_HRSVA` lists five “Cold-passage attenuated” variants (positions 102, 218, 379, 447, and 523), none of which were retained by our procedure. So does `NCAP_HRSVA` (V267I). Since the method of attenuation (and indeed of vaccine production) is usually not known for our vaccine genomes, it is impossible to use these known mutations as positive controls, and we can only note that the mutations we identified are hitherto unknown.

### Affected Subtypes

Our procedure only identified mutations in vaccines derived from HRSV subtype A, but the failure to find mutations in type B is possibly due to the lower number of sequences available from that subtype, as this would lower the power of the statistical test used to identify positions with distinct allele frequency distributions between wild-type and vaccine sequences. Another possible explanation would be that vaccines only target subtype A.

### Limitation to Coding Regions

Our procedure is limited to coding regions of the HRSV genome. Any mutation falling outside of coding regions will thus be missed, and attenuating mutations in non-coding regions have been reported (in transcription start sites, for example^29^). However, of the six attenuating point mutations reported by Karron *et al.*^12^, five were in coding regions, which suggests that limiting our procedure to coding regions is unlikely to cause it to overlook all (or perhaps even most) mutations of interest.

### Incompatible Mutations

While the design of vaccines by reverse genetics will likely benefit from an expanded list of attenuation-linked mutation candidates, it should be borne in mind that mutations may be pairwise incompatible.^30^

## Conclusion

We identified a set of positions, in the genome of the human orthopneumovirus subtype A (HRSV-A), which exhibit significantly different allele frequencies sequences annotated as “vaccine” with respect to sequences not so annotated. This set is likely to contain any position under positive selection pressure in the process of attenuation carried out in the production of candidate live HRSV vaccines.

We also found that the vaccine sequences almost never have the wild-type allele at any of these positions. If, as seems likely, the vaccine sequences are not closely related, this means that the separate attenuation processes have independently converged on the same mutations.

We found no comparable mutations in subtype B, but this can be due to small sample size rather than lack of selection pressure.

These findings adds to the current understanding of the genome-scale effects of attenuation in HRSV, and may find application in the direct engineering of live attenuated vaccines.

## Methods

### Overview

We aligned homologous gene sequences from vaccine candidates and wild-type viruses together (but keeping the two subtypes separate), and searched for positions with significantly different allele frequencies in the two groups. To identify positions with different allele frequencies in vaccine vs. wild-type sequences, we first had to distinguish these two categories, and to classify sequences by gene and subtype when not annotated. The detailed procedure (implemented using Python^31^ (including BioPython^32^), Bash^33^, SQLite^34^, and R^35^) follows. A workflow diagram of the whole procedure is shown in supplemetary figure S1 (PDF and SVG).

### Download and Sorting of Raw Sequence Data

We downloaded all entries with key phrase `Respiratory syncytial virus` from GenBank^36^. There were 28,248 at the time of download - see supplementary file `all.gb`. This included both whole-genome and partial HRSV sequences, of both subtypes, mostly wild type but with a small minority annotated as “vaccines”. Although it would be more accurate to refer to these as “vaccine candidates”, this phrase is unwieldy and we will refer to them as “vaccines” in the rest of this article.

A small number of entries were not from human orthopneumovirus, and were discarded.

We then separated wild-type from vaccine sequences based on annotation: any entry containing the string `VACCINE`, `ATTENUAT`, or `PATENT` in the Definition, Source, Reference Titles, or Comments fields of the GenBank entry was considered a vaccine sequence (this assumes that patented sequences are vaccines); all others were considered wild-type.

Likewise, we separated the two subtypes of wild-type viruses by annotation, using various regular expression searches for patterns such as `type (A|B)` in the Source, Organism, or Definition fields of the Genbank entry, as well as in its Feature Table (for the full details, see the source code in file `allRSVtoCSV.py` in the supplementary data). Entries whose subtype could not be determined were discarded. Vaccine sequences were rarely annotated for subtype, so they were not included in this step, indeed the results from this step were used to determine the subtype of vaccine sequences (see below).

We then used entry annotations to extract the coding sequences of each wild-type entry of recognizable subtype: we scanned the GenBank entry’s Feature Table for coding sequences (features of type `CDS`), within which we searched for keys such as `gene` or `product`. We then folded the resulting names into a controlled name list (e.g., entry annotations “L”, “l”, “polymerase”, and “large polymerase”, which in fact denote the same gene, were all mapped to “L”). The complete mapping is available as `gene_name_map.py` in the supplementary data. As was the case for subtype, vaccine sequences were rarely annotated for gene, so they were not included in this step either, but the results from this step were also used to determine the gene of vaccine sequences (see below).

We rejected any wild-type sequence containing unknown nucleotides (`N`)s. We also discarded partial gene sequences by retaining only those sequences that were at least 9/10 of the maximum length for the corresponding gene and subtype. The rationale for this is that sequences of poor quality or insufficient length can not only slow down the alignment process while contributing little information, but can also cause too many columns to be dropped when building phylogenetic trees.

The procedure up to this point yielded i) twenty sets of wild-type DNA sequences of recognized subtype and gene (namely, ten genes each for both subtypes), all devoid of `N`s, and full-length or nearly so; and ii) a set of (as yet) uncharacterised vaccine sequences.

### Classifying Vaccine Sequences

Since the vaccine sequences rarely featured annotation of gene or subtype, these were inferred by searching them for matches of profile hidden Markov models (HMMs) built from wild-type sequences of known gene and subtype, as follows.

#### Reference (Wild-type) HMMs

All twenty wild-type sequence sets obtained as described above were aligned using Mafft^37^, with default parameters. The resulting alignments were input to the HMMER^38^ 3.1 package’s `hmmbuild` program, also using default parameters, yielding one model for each (subtype, gene) combination. The models were collected into a database using HMMer’s `hmmpress`.

All vaccine sequences were then scanned, using HMMer’s `nhmmscan`, for matches of any HMM in the database (it being understood that a given vaccine sequence may match more than one HMM, albeit at different, non-overlapping positions - as is expected, for instance, of whole-genome vaccine sequences). Matches were classified (i.e., attributed to a subtype and gene) according to the highest HMM score, and sorted into separate files by subtype and gene.

### Selecting Vaccine Sequences

We computed a phylogenetic tree for each gene, as follows: first, we pooled the wild-type sequences that were used for alignment (namely, those that matched all our quality criteria) of the two subtypes, yielding a single sequence set per gene. Vaccine sequences were added to the corresponding set, subject to the condition that they also be at least 90% as long as the maximum length for the corresponding gene - again, in order to minimize the loss of aligned positions when computing the tree. Finally, the sequence of the orthologous gene from the bovine orthopneumovirus (`NC_0 0198 9`) was added to each set as an outgroup.

Each of the ten resulting sets was aligned with Mafft, using default parameters. The resulting FastA was converted to PHYLIP^39^ format using the EMBOSS^40^ package’s `seqret` program. Trees were computed with Phyml^41^ version 3.0, using the general-time-reversible-Gamma model, four substitution rate categories, and maximum-likelihood estimates of base frequencies, transition/transversion ratio, as well as gamma distribution shape parameter. The resulting trees, as well as various graphical representations of them, are available in the supplementary data.

#### Reduction of Similarity by Inheritance: Selection of Clade Representatives

We selected one vaccine sequence from each purely- or almost-purely-vaccine, completely unbalanced clade, as revealed by the phylogenetic trees. These clade representatives were randomly drawn from the two sequences with the largest number of ancestors within the clade (see Discussion for rationale).

#### Aligning the Vaccines to the Wild-type References

The clade representatives were added to the corresponding alignments of wild-type sequences using Mafft with the `—add` option. This aligns the additional (namely, vaccine) sequences without changing the original (i.e., wild-type) alignment.

#### Frameshifts

Examining the sequences with an alignment viewer^42^ showed that twelve sequences had frame shift mutations. These sequences were removed and the alignments recomputed until no frame shifts were observed.

#### Finding Mutated Positions

The search for positions likely to be under positive selection pressure from the attenuation process was carried out using a Python script (see `find_mutations.py` in the supplementary data) written for this purpose, its main steps are as follows:

- for each position in the alignment:
  1. identify the most frequent allele (pooling vaccine and wild-type sequences), define this to be the *major allele*;
  2. fold the remaining three alleles into category *minor allele;*
  3. compare the frequency of the major versus minor alleles, in wild-type versus vaccine sequences, using Fisher’s exact test;
  4. if the test’s *p*-value is below some predefined threshold, register this position as a vaccine-linked mutation position, then determine if the most frequent allele in vaccine versus wild-type sequences result in different amino acids: if so, report the position as a vaccine-linked, non-silent mutation position.

## Data Availability

Code, data, supplementary figures and the main intermediary results are available from https://figshare.com/articles/dataset/2020_RSV_article/13102796, doi:10.6084/m9.figshare.13102796.

## Author contributions statement

J.F. and T.J. designed the experiments, except for the statistical aspects which were handled by N.C. T.J. wrote and ran the code implementing the experiments, and wrote the manuscript. L.K. and T.H. contributed expertise in virology and to the experimental design. All authors reviewed the manuscript.

## Additional information

The authors declare no conflict of interest. No funding was sought or received.

