## Supplementary material for "Genomics of Human Respiratory Syncytial Virus Vaccine Attenuation": figure S1

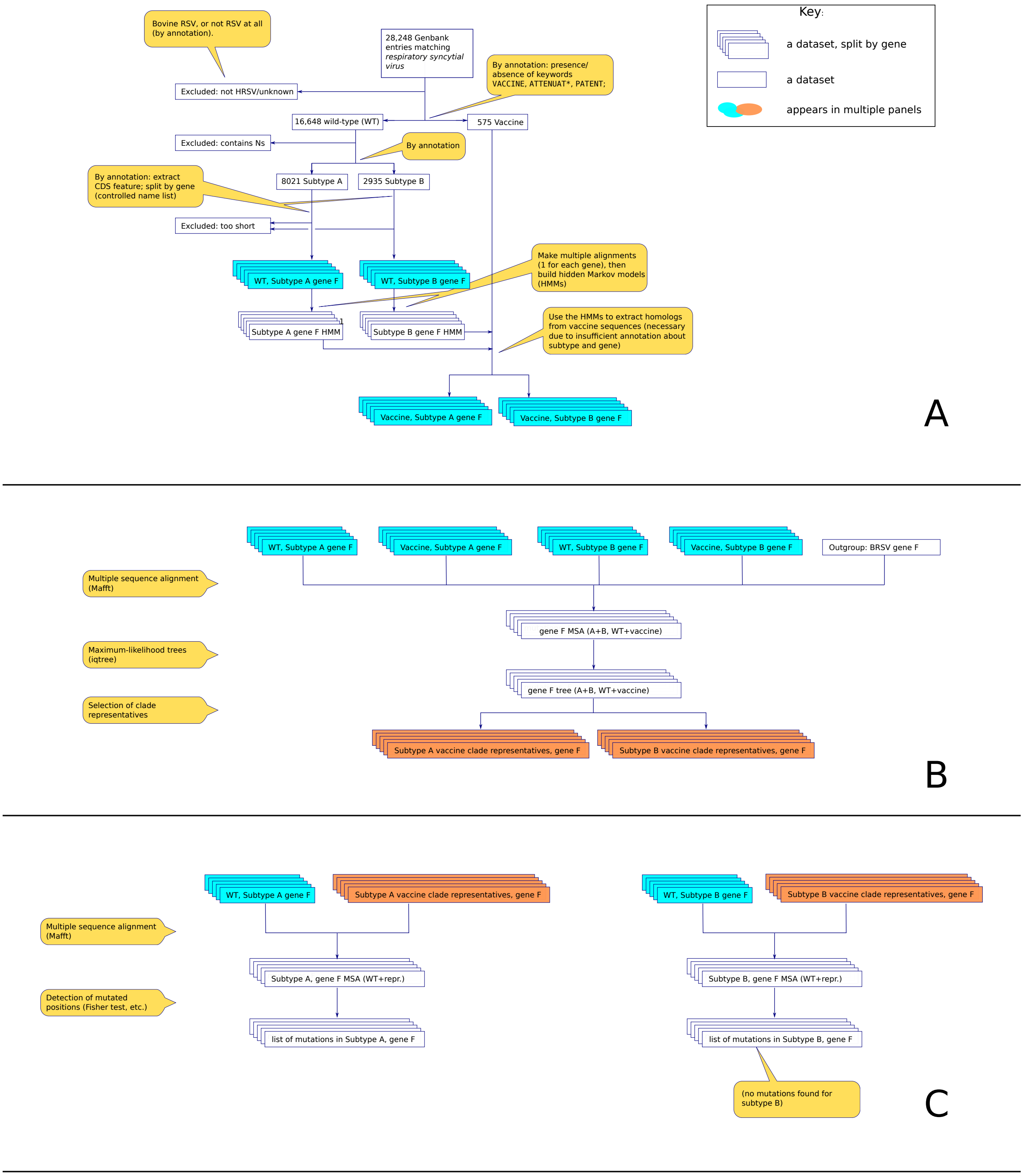


**Data Flow:**

**A**: Extracting and sorting sequences by gene, subtype and origin (WT/vaccine). Input is the set of GenBank entries pertaining to RSV. Outputs are sets of sequences, one each for each gene/subtype/origin combination (cyan). Classification is by annotation (WT) or by sequence similarity to WT (vaccines).
**B**: Building phylogenies and extracting clade representatives. Inputs are the sequence sets from step **A**, outputs are sets of sequences representative of each clade (orange). See text for rationale.
**C**: Finding mutated positions. Inputs are results from steps **A** and **B**, outputs are lists of positions which have significantly different major/minor allele frequencies among vaccines by comparison with WT sequences. Significance is measured by Fisher's exact test.
